# Observation of E-cadherin Adherens Junction Dynamics with Metal-Induced Energy Transfer Imaging and Spectroscopy

**DOI:** 10.1101/2023.12.16.571977

**Authors:** Tao Chen, Narain Karedla, Jörg Enderlein

## Abstract

Epithelial cadherin (E-cad) mediated cell-cell junctions play a crucial role in the establishment and maintenance of tissues and organs. In this study, we employed metal-induced energy transfer imaging and spectroscopy to investigate variations in intermembrane distance during adhesion between two model membranes adorned with E-cad. By correlating the measured intermembrane distances with the distinct E-cad junction states, as determined by their crystal structures, we probed the dynamic behavior and diversity of E-cad junctions across different binding pathways.

Our observations led to the identification of a transient intermediate state referred to as the X-dimeric state and enabled a detailed analysis of its kinetics. We discovered that the formation of the X-dimer leads to significant membrane displacement, subsequently impacting the formation of other X-dimers. These direct experimental insights into the subtle dynamics of E-cad-modified membranes and the resultant changes in intermembrane distance provide novel perspectives on the assembly of E-cad junctions between cells. This knowledge enhances our comprehension of tissue and organ development and may serve as a foundation for the development of innovative therapeutic strategies for diseases linked to cell-cell adhesion abnormalities.

**Significance Statement:** In this study, we employed metal-induced energy transfer (MIET) imaging and spectroscopy to track variations in intermembrane distance during the adhesion of two membranes mediated by epithelial cadherin. Leveraging the high spatial resolution of MIET, we explored the dynamics of cadherins across various binding pathways. Furthermore, we successfully captured a transient intermediate state known as the X-dimer and revealed its ability to communicate with other X-dimers through membrane displacement. These discoveries offer valuable mechanistic insights into the dynamics of cadherin junctions.

---

Epithelial cadherin (E-cad) is a prominent member of the classical cadherin family, categorized as a type I cadherin. It serves a crucial role in facilitating calcium-dependent adhesion among epithelial cells, thereby contributing to the formation of adherens junctions on the cell surface (1–5). By mediating the adhesion between epithelial cells, E-cad contributes to the establishment and maintenance of cellular interactions within tissues. The formation of adherens junctions, enabled by E-cad’s activity, promotes the cohesion and structural stability necessary for the proper functioning of organs and tissues.

However, the regulation of E-cad expression and activity is not static but rather subject to dynamic changes. These alterations play a vital role in various physiological and pathological processes (6). For instance, the loss or downregulation of E-cad expression is often observed in metastatic cancers, where it contributes to the acquisition of invasive and migratory properties by cancer cells (7). In such cases, E-cad’s normal adhesive function is disrupted, allowing cancer cells to detach from their original location and invade surrounding tissues.

Additionally, during inflammatory responses, the dysregulation of adherens junctions and E-cad expression can compromise the integrity of epithelial barriers. This disruption can result in increased permeability and allow the infiltration of harmful substances, exacerbating the inflammatory process. Thus, the dynamic regulation of E-cad in these scenarios becomes crucial for maintaining tissue homeostasis and preventing the progression of inflammatory disorders (8–12).

The classic form of E-cad is comprised of five structurally similar extracellular cadherin-like domains (EC domains) arranged in tandem. These domains are connected by conserved calcium-binding linker regions. E-cad also possesses a single transmembrane segment and a highly conserved cytoplasmic tail. Numerous studies have underscored the pivotal role of the EC domains of cadherin in regulating the assembly of cell-cell junctions (13–16).

The process of cell-cell adhesion involves multiple steps, encompassing the dynamic transition between various molecular states, ranging from free monomers to trans-dimers, clusters of E-cad molecules, and interactions with intracellular domains (Fig. 1a) (17). A weak and transient X-dimer has been reported to exist during the adhesion process, acting as a kinetic intermediate that lowers the energy barrier during the formation of st rand-swapped S-dimers in adherens junctions (13). Initially, this X-dimer was identified through sit e-directed mutations in E-cad (18, 19). More recently, the X-dimer was finally observed in wild-type cadherins using cryo-electron microscopy (20).

**Fig. 1.**
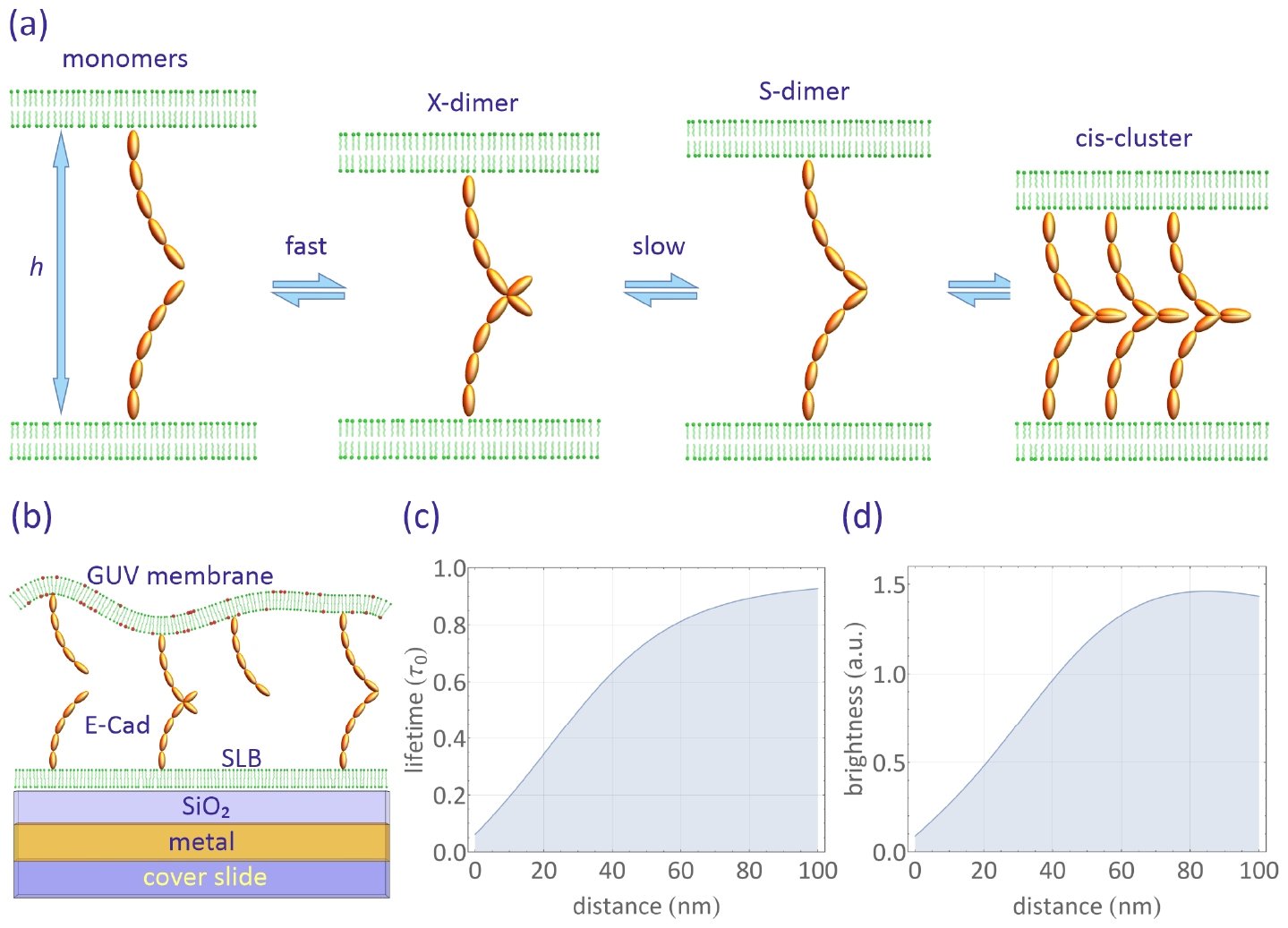
(a) Schematic representation of classic E-cad states during clustering: This diagram illustrates the various states classic E-cads can assume during clustering. The process of trans-dimerization, where E-cads join together, is mediated by the extracellular subunits EC1 and EC2. It involves at least two distinct structural states: initially, a short-lived X-dimer forms from individual monomers, which then transition into a more stable S-dimer. The S-dimer has the ability to cluster further by establishing lateral interactions within the same cell, known as *cis*-interactions, mediated by EC1-EC2 connections to the actin cytoskeleton. These *cis*-interactions between E-cads on the same cell form a lattice-like structure as proposed in adherens junctions. Importantly, these transitions between molecular states can also occur in reverse, involving the disassembly of *cis*-interactions and transitions from S-dimers back to X-dimers and monomers. (b) Experimental Configuration: This schematic illustrates the setup used in our experiment. E-cad proteins are positioned between a supported lipid bilayer (SLB) on a gold film, separated by a silica spacer, and a giant unilamellar vesicle (GUV) that is highly labeled with fluorescent dye. (c, d) Calculated fluorescence (c) lifetime and (d) intensity dependence on distance: This graph displays the calculated relationship between fluorescence lifetime (expressed as τ / τ _0_, with τ _0_ representing lifetime in free space) and fluorescence intensity relative to the distance above the substrate surface. The curves are calculated for a dipole with an emission wavelength of 690 nm and a random orientation. The substrate used for these calculations consists of multiple layers, including 10 nm silica, 1 nm titanium, 10 nm gold, 2 nm titanium, and a glass coverslip, arranged from top to bottom.

E-cads have been the subject of extensive investigation utilizing various experimental and theoretical approaches, which have significantly contributed to our understanding of E-cad-mediated intermembrane adhesion (13–15, 17, 21–27). For example, NMR relaxation dispersion spectroscopy has enabled the measurement of kinetic parameters in the trans-dimerization of E-cad EC fragments in solution (13). Single-molecule Förster resonance energy transfer (smFRET) studies have provided evidence of the mutual cooperativity of cis/ trans interactions (14, 22). Fluorescence microscopy has revealed the involvement of a nucleation process in junction formation (24, 28), while reflection interference contrast microscopy (RICM) and computational simulations have shed light on the role of membrane fluctuations in mediating the interaction between E-cad bonds (23).

However, limited attention has been given to variations in the EC5-EC5 distance during adhesion. Crystal lattice studies have identified EC5-EC5 distances of 29 nm for the X-dimer, 37 nm for the S-dimer, and 19 nm for the cis-cluster of E-cad (20, 29, 30). These changes in EC5-EC5 distance offer an alternative means of monitoring E-cad adhesion; however, resolving such minute distance changes requires a high-resolution technique.

Metal-induced energy transfer (MIET) imaging and spectroscopy represent s a cutting-edge super-resolution fluorescence technique ideally suited for such inquiries (31–33). MIET excels at precisely localizing fluorophores in the axial dimension with nanometer-scale accuracy, employing conventional fluorescence lifetime imaging microscopy. This technique capit alizes on the electrodynamic near-field coupling between an excited fluorophore and surface plasmons within a planar metal film.

The distance-dependent energy transfer from the fluorophore (donor) to the metal (acceptor) in MIET leads to a distinctive modulation of both fluorescence lifetime and brightness. These changes are intimately linked to the fluorophore’s proximity to the metal surface. By meticulously measuring fluorescence lifetimes, we can accurately determine the distance between the fluorophore and the metal, employing a suitable theoretical model.

We harnessed the power of MIET to dissect the intermembrane adhesion process mediated by E-cads. Our focus lies in exploring the variations in membrane height, particularly the EC5-EC5 distance, during the adhesion of two E-cad-modified membranes. By employing this innovative approach, we can unravel the dynamic landscape of E-cad binding states, illuminating the intricacies of the adhesion process. The utilization of MIET grants us the capability to gain insights into the nanoscale alterations occurring in E-cad-mediated intermembrane adhesion. This, in turn, provides information regarding the behavior and interactions of E-cads during the intricate process of cell-cell adhesion.

## Results

We used a biomimetic system to mimic cad-mediated cell adhesion, observing adhesion events between partially fluid supported lipid bilayers (SLBs) and giant unilamellar vesicles (GUVs). The SLBs were prepared using 1,2-dipalmitoyl-sn-glycero-3-phosphocholine (DPPC) doped with 1 mole% acyl chain-labeled 1-acyl-2-[12-[(7-nit ro-2-1,3-benzoxadiazol-4-yl)amino]dodecanoyl]-sn-glycero-3-phosphocholine (NBD-PC), 5 mole% 1,2-dioleoyl-sn-glycero-3-[(N-(5-amino-1carboxypentyl)iminodiacetic acid)succinyl] nickel salt (Ni-NTA-DOGS), and 1,2-dioleoyl-sn-glycero-3-phosphoethanolamine-N-[methoxy(polyethylene glycol)-2000](DOPE-PEG2000). The GUVs were prepared using 1-stearoyl-2-oleoyl-sn-glycero-3-phosphocholine doped with 5 mol% NTA-DOGS, 1 mol% DOPE-PEG(2000), and 1 mol%Atto655 headgroup-labelled 1,2-Dipalmitoyl-sn-glycero-3-phosphoethanolamine (Atto655-DPPE). These membranes were decorated with E-cad extracellular (EC) domains (Fig. 1b).

The E-cad receptor was in the form of a dimeric chimera, with all five E-cad EC domains genetically fused to an immunoglobulin Fc-fragment exhibiting a hexahistidine t ag (his6). The SLBs were prepared using the Langmuir-Blodgett Langmuir-Schäfer technique on a substrate consisting of a 10 nm Au film and a 10 nm silica spacer. The distal leaflet of the SLB was doped with the Ni-nitrilotriacetic acid (NTA) chelating lipid, which can further bind to the his6 tags of the E-cad. The GUVs were also functionalized with E-cad EC at a concentration of 5 mol%. To induce membrane fluctuations and excess area in the vesicles, we created an osmolarity difference between the inside of the GUVs (230 mOsm l^− 1^, sucrose solution) and the outer buffer (300 mOsm l^− 1^, phosphatebuffered saline, PBS, mixed with 0.75 mM CaCl_2_ solution). No adhesion process was observed in the absence of a osmolarity difference (Supporting Information, Fig. S2).

A high dye concentration (1 mol%, Atto655-DPPE) was used to eliminate the contributions form lateral diffusion to the observed fluorescence fluctuations: At such a high concentration, fluorescence intensity fluctuations arising from dye diffusion in and out of the confocal volume are negligible (Supporting Information, Figure S1). The adhesion process was monitored using MIET imaging/ spectroscopy. Figure 1c shows the calculated dependence of lifetime and brightness as a function of height from the silica surface (see Materials and Methods). This allowed us to translate a measured lifetime into an axial distance value using the lifetime-distance calibration curve. In this study, only the membrane of the GUV was labeled, and its height over the surface was precisely determined from the lifetime values of the labeling fluorophores.

The fluorescence imaging and lifetime measurements for the samples were conducted using confocal scanning microscopy, employing pulsed laser excitation and time-correlated single-photon counting (TCSPC) for fluorescence lifetime measurements, as detailed in the Materials and Methods section. Initially, we scanned the samples to identify suitable GUVs, including only those with a diameter exceeding 15 μm for further analysis.

In Fig. 2a, one can observe a fluorescence FLIM image of GUVs containing E-cad on the SLBs after 1 hour of incubation. The FLIM image reveals two distinct binding states in the proximal membranes of the GUVs:

**Fig. 2.**
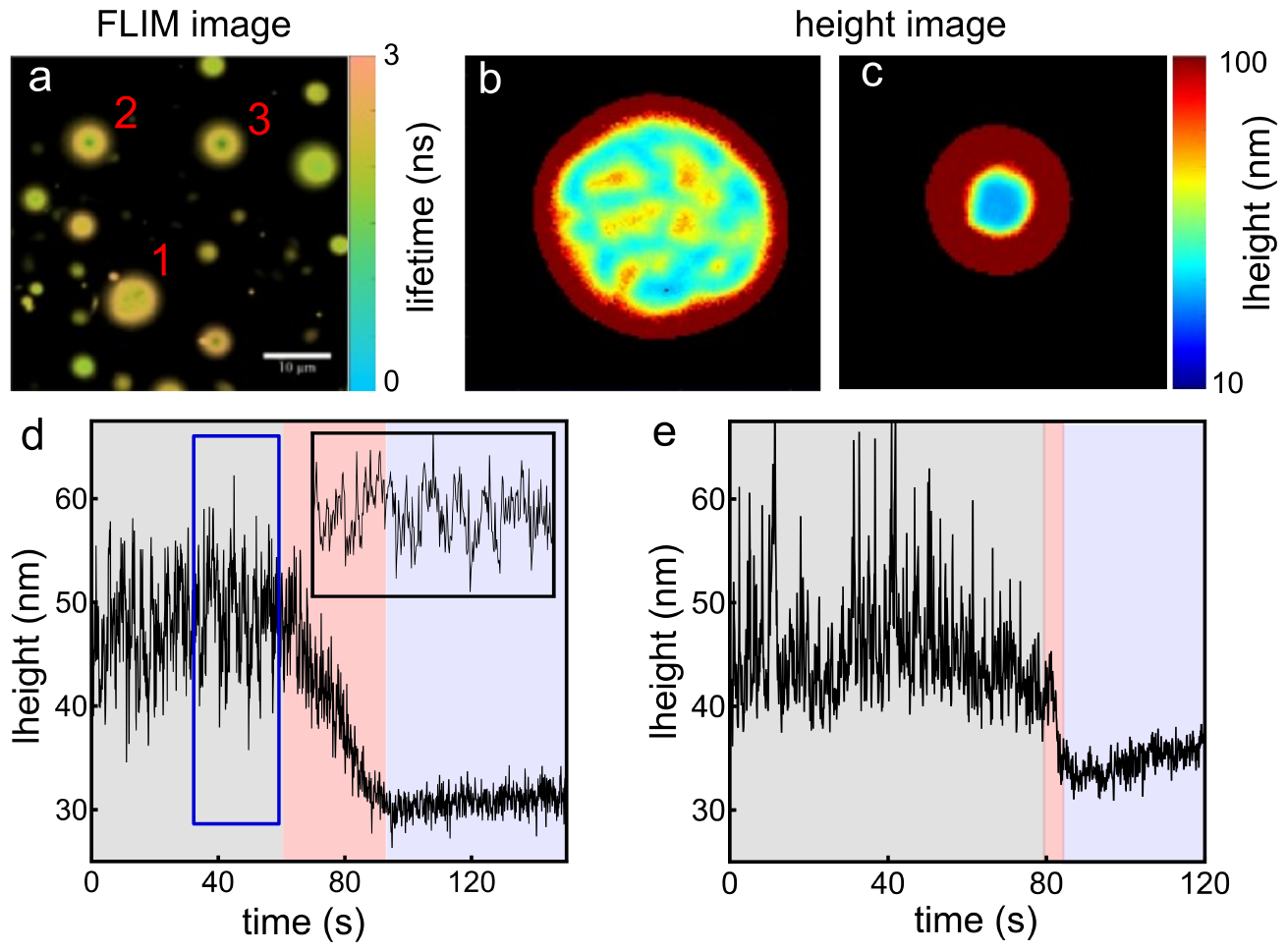
Figure 2. (a) FLIM image of fluorescently-labeled GUVs containing E-cad on SLBs. The image was measured 1 h after incubation of the GUVs within the chamber. (b,c) Reconstructed height images for GUVs in different adhesion states. (d,e) Height traces showing height variations during E-cad clustering for measurements on a partially fluid SLB (d), and for a fluid SLB (e). Inset at top-rightin (d): Zoom-in of the trace marked by the blue rectangle.

- The lifetimes and contact areas of the proximal mem-branes for GUV-2 and GUV-3 are significantly shorter and smaller than those of GUV-1.
- The lifetime distribution for the contact area is uniform for GUV-2 and GUV-3, while it displays more heterogeneity for GUV-1.

Utilizing the MIET calibration curve depicted in Figure 1c, we transformed the FLIM images into height images. These height images revealed that the average bottom height of GUV-2 and GUV-3 above the surface is approximately 35 nm, while GUV-1 exhibits a greater height of 56 nm. Factoring in the thickness of the SLBs (approximately 4 nm) and the hydration layer (about 2 nm) (32), the size of the Fc fragment (d_Fc_ = 7 nm, as determined from the crystal structure) (34), and the tile angle of the E-cad in the clustered state (approximately 30°, determined from the crystal structure) (26, 29), we calculated that the EC5-EC5 distance for GUV-2 and GUV-3 is approximately 20 nm, whereas for GUV-1, it measures roughly 35 nm.

This analysis provides confirmation that GUV-2 and GUV-3 have indeed formed the *cis*-cluster junctions, as the calculated 20 nm EC5-EC5 distance closely aligns with the crystal structure of a *cis*-cluster (19 nm). However, due to the observed inhomogeneity in GUV-1, classifying the binding state for GUV-1 solely based on its average height proves challenging. This observation implies that, in addition to the E-cads on the surface of the membranes (both GUV and SLB) that culminate in the final *cis*-clustered state (as seen in GUV-1 and GUV-2), some E-cads do not contribute to the formation of the ultimate *cis*-cluster. In fact, during our 2-hour observation period, membranes such as GUV-1 consistently maintained this heterogeneous pattern. This observation aligns with a previous report indicating the existence of a nucleation process during E-cad junction formation, resulting in an all-or-nothing feature (24).

In the initial phase of *cis*-cluster interface formation, we meticulously monitored the E-cad-mediated adhesion process via FLIM imaging, see Movie S2. Initially, the giant unilamellar vesicles (GUVs) positioned themselves above the supported lipid bilayers (SLBs) to establish a contact zone. This contact zone subsequently contracted during the adhesion process due to the robust enrichment of the *cis*-cluster. To track the height variations throughout this process, we initiated point measurements to record fluorescence intensity traces (as detailed in Supporting Information, Figure S4).

Subsequently, we transformed these fluorescence intensity traces into a height traces by binning them at a 100 ms resolution. For each time bin (containing between 1 × 10^4^ and 5 × 10^4^ photons), we constructed a fluorescence decay histogram and determined the fluorescence lifetime using a multi-exponential decay model. These obtained lifetimes were then converted into height values utilizing the calibration curve presented in Fig. 1c.

A typical height trace representing one GUV adhesion event is illustrated in Fig. 2d, which can be segmented into three distinct phases. The initial black region in Fig. 2d signifies the waiting period preceding E-cad clustering. The subsequent red segment corresponds to the clustering process it self, and the subsequent blue area represents the time trace of E-cad after clustering.

During the course of the curve depicted in Fig. 2d, the membrane Initially exhibited significant height fluctuations within the range of 45-55 nm (black region). Subsequently, it gradually decreased to a lower height of approximately 31 nm with reduced fluctuations (blue region). The initial height range of 45-55 nm corresponds to an EC5-EC5 distance of 23-33 nm, assuming no tilt of the E-cad. In contrast, the final height of approximately 31 nm corresponds to an EC5-EC5 distance of 16 nm, assuming a tilt angle of 30 degrees for the E-cad.

Taking into account crystal structure information on E-cads, the EC5-EC5 distance range of 23-33 nm suggests that the initial state of the E-cad in our measurement is a dimeric state (S-/ X-dimer), while the final state of 16 nm represents the ultimate *cis*-clustered state. It is worth noting that we do not observe the process of monomer to dimer formation due to the ultrafast dimerization rate (3.8 × 10^4^ *s*^− 1^) of E-cad monomers. However, we cannot assume that monomers do not exist in the initial state because E-cads dynamically transition between different states.

In Fig. 2d, the red segment encompasses the timescale of the *cis*-clustering process. We discovered that this process strongly depends on the mobility of the membrane. For instance, when we replaced the partial fluid SLB (composed of DPPC+ NBD-PC) with a fluid SLB made of pure SOPC lipid, the clustering rate increased by at least an order of magnitude (as shown in Fig. 2e, and further detailed in Supporting Information, Fig. S4).

In instances where cluster formation did not occur, the lower membrane of the giant unilamellar vesicle (GUV) exhibited continuous fluctuations at a consistent height, as visually demonstrated in Supporting Movie S2. In our MIET measurements, the GUV membrane fluctuates above the gold-coated surface, causing variations in fluorescence intensity (as shown in Fig. 1c). Importantly, we verified that these intensity fluctuations were the result of vertical displacements of the membrane and not caused by the lateral diffusion of individual fluorophores. This is due to the high fluorophore label concent ration (1 mol%), which excluded any contribution from the lateral diffusion of the fluorophores to the observed intensity fluctuations (Supporting Information).

Furthermore, we conducted point measurements on this fluctuating membrane and analyzed the time trace of fluorescence intensity. Surprisingly, we discovered that the intensity distribution exhibited asymmetry, and numerous ‘dip’ signals were evident in the intensity-time traces (as depicted in Fig. 3a, and detailed in Supporting Information, Fig. S5). As a control experiment, we did not observe such ‘dip’ signals in the GUV membrane lacking E-cad or in the membrane after the formation of the *cis*-cluster (Fig. 3b).

**Fig. 3.**
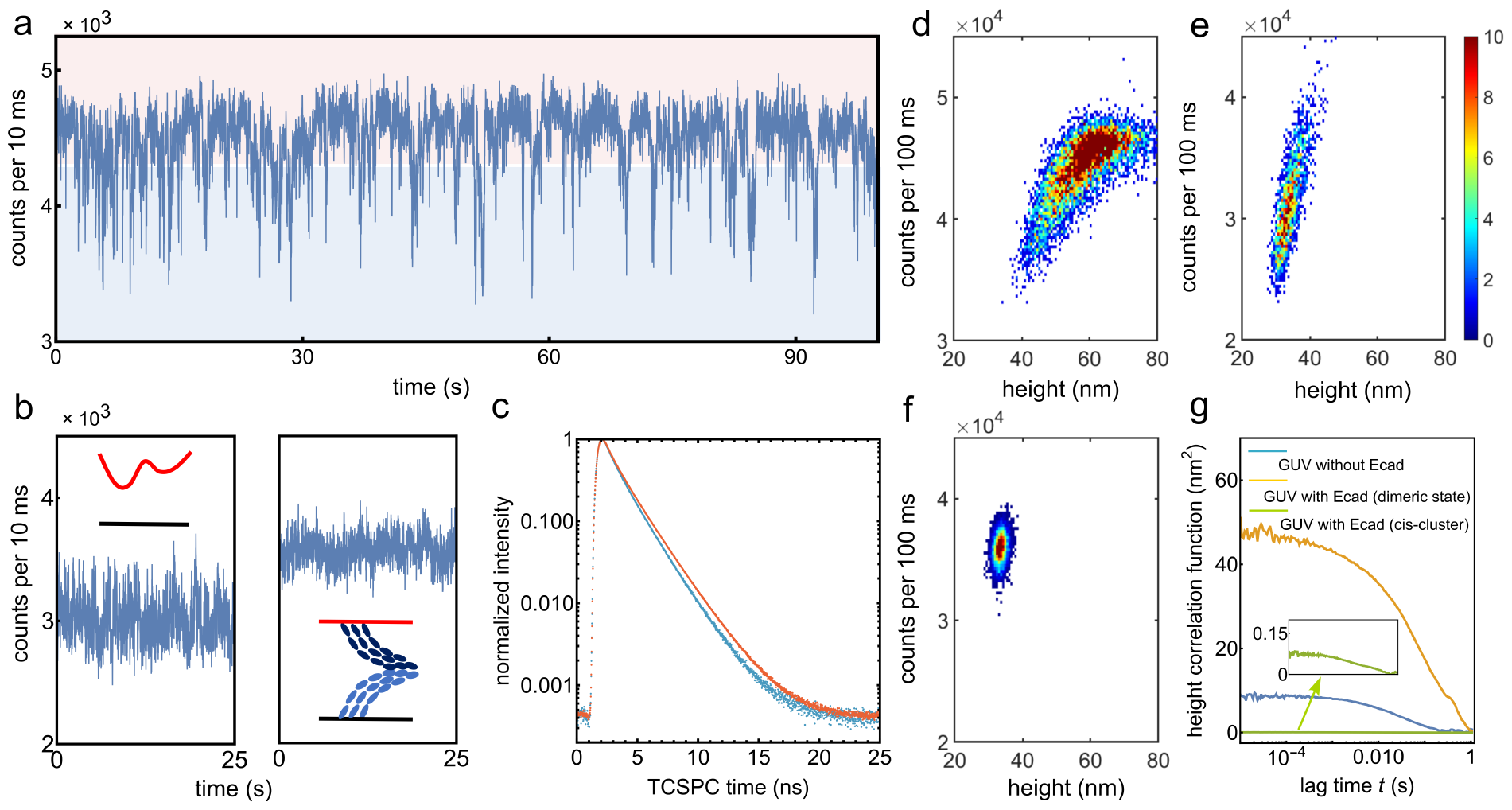
(a) A typical fluorescence intensity trace recorded from a proximal membrane of highly labeled GUV with E-cad on an E-cad-modified SLB. (b) Fluorescence intensity traces recorded from the GUV-SLB system without any protein (left), and from the membranes formed the *cis*-clusters (right). (c) Fluorescence lifetime decay curves for the high state (red curve) and low state (blue curve), derived from the trace in panel (a). (d,e,f) Two-dimensional histograms of the fluorescence intensity and the height from the surface for (d) the membrane with E-cads, (e) the membrane without any proteins, (f) the membranes formed the final *cis*-clusters. (g) Height autocorrelation functions for a membrane without E-cad (blue), a membrane with E-cad but not forming the *cis*-clustered state (yellow curve), and a membrane forming the *cis*-clustered state (green curve). Inset shows the enlarged plot of the green curves.

Returning to the first case that led to the formation of the final *cis*-cluster, we also observed these ‘dip’ signals in certain time traces within the waiting time interval preceding E-cad clustering (Fig. 2e, inset).

In our quest to unravel the origin of the intriguing ‘dip signals,’ we employed a two-threshold method to demarcate the ‘dip signal’ (referred to as the low state) from the baseline (referred to as the high state) within the intensity-time trace (see Supporting Information). Subsequently, we constructed fluorescence lifetime decay histograms for these two distinct states and determined their respective fluorescence lifetimes (as illustrated in Fig. 3c).

Through fitting the fluorescence decay curves using a multiexponential model, we extracted lifetimes of 1.65 ns for the high state and 1.51 ns for the low state, respectively. These lifetimes corresponded to heights of 55 nm and 47 nm above the silica surface. Notably, the heights of 55 nm and 47 nm correlate with EC5-EC5 distances of 33 nm and 25 nm, aligning remarkably well with the crystal structure values of the S-dimer (37 nm) and X-dimer (29 nm) configurations of E-cad. Furthermore, the brief duration of the low state in the intensity-time trace corresponds to the short-lived and unstable structure of the X-dimer. Thus, we attribute the origins of the high state and low state to the membrane positions associated with the S-dimer and X-dimer of E-cad configurations, respectively.

Moreover, we conducted experiments to validate that the consecutive ‘dip signals’ emanate from the continuous formation of X-dimers within the confocal volume, rather than from the lateral diffusion of X-dimers. This confirms that X-dimers formed within the confocal volume and did not undergo significant diffusion in and out of the confocal volume (details provided in the Supporting Information).

It is essential to note that E-cad-mediated intermembrane adhesion is a multifaceted process with various pathways, including monomer-involved routes. These pathways encompass interactions such as monomer binding to S-/ X-dimers, backward unbinding from S-/ X-dimers to monomers, and *cis* binding/ unbinding of monomers. Unfortunately, in our measurement, we cannot distinguish these specific monomer-involved processes. This limitation arises from the fact that our measurement relies on monitoring membrane height (or EC5-EC5 distance), and monomers exist in a free state without a distinct height signature. Consequently, changes in monomer populations do not induce discernible alterations in membrane height.

For instance, consider the scenario of backward unbinding from an X-dimer to monomers. Given that a significant majority of E-cads are in an S-dimeric state, even if an X-dimer dissociates into monomers within each membrane, the membrane’s height would still primarily reflect that of an S-dimer. Therefore, we cannot definitively conclude that the abrupt transition from the low state to the high state in the time trace exclusively represents the process of converting X-dimers to S-dimers. This transition encompasses a combination of two potential pathways: X-dimer to S-dimer conversion and X-dimer to monomer conversion.

We further delved into the membrane fluctuations or displacements induced by the formation of X-dimers. To achieve this, we computed membrane heights for a binning time of 100 ms from the intensity trace and conducted an analysis that considered both fluorescence intensity and membrane height fluctuations (as computed from the fluorescence lifetime) throughout the measurement process. As illustrated in Fig. 3d, a two-dimensional (2D) histogram reveals correlations between emission intensity and membrane height for the E-cad-modified membranes. This histogram depicts a positive correlation between fluorescence intensity and membrane height, consistent with the theoretical intensity-height curve presented in Fig. 1c.

The height distribution spans a wide range, varying from 40 nm to 75 nm, with a predominant concentration around ∼ 60 nm. In contrast, membranes lacking E-cad exhibit a narrower height range, spanning from 30 nm to 45 nm. Moreover, for membranes in the *cis*-clustered state, the height distribution is even more confined, ranging from 32 nm to 36 nm. Notably, the formation of X-dimers leads to heightened membrane fluctuations compared to the other two control membranes in the experiment (Fig. 3e, f).

This increased fluctuation arises from a combination of two contributing factors. Firstly, self-fluctuation occurs as a result of osmolarity differences. Secondly, dynamic structural changes within the dimers contribute significantly to this enhanced fluctuation.

To quantitatively assess membrane fluctuation, we constructed the membrane displacement autocorrelation function (g_h_ (t)) based on the fluorescence intensity autocorrelation function (g_I_ (t)). The g_I_ (t) is derived from the fluctuations in fluorescence intensity within the confocal volume. As demonstrated earlier, the fluctuations in fluorescence intensity exclusively originate from the physical displacement of the membrane. Consequently, we can convert the *g*_*I*_ (*t*) to *g*_*h*_ (*t*) by utilizing the relationship between fluorescence intensity and membrane height (see Materials and Methods). The *g*_*h*_ (*t*) is defined as *g*_*I*_ (*t*) = < *δh*(0)*δh*(*t*) >, where *δh*(*t*) represents the instantaneous membrane displacement . The height correlatione function provides the root mean square displacement 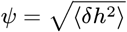 of the membrane, as well as the relaxation time *τ* ^∗^, defined as the time point at which *g*_*h*_ (*t*) has declined to half of its maximum value (35).

As illustrated in Fig. 3g, the displacement amplitude *ψ* for a GUV membrane with E-cad is 6.9 nm, whereas it is *ψ* = 2.9 nm for the membrane lacking E-cad. In the latter case, membrane fluctuations are solely driven by differences in osmolarity. Following the formation of the final *cis*-clusters, membrane fluctuations become negligible and fall below the detection threshold (Fig. 4g). Furthermore, the relaxation time of the membrane fluctuations for GUVs with E-cad (*τ* ^∗^ =0.1 s) is notably slower than that for GUVs without E-cad(*τ* ^∗^ = 0.04 s)This difference is attributed to the slower structural interconversion between S- and X-dimers in the presence of E-cad.

**Fig. 4.**
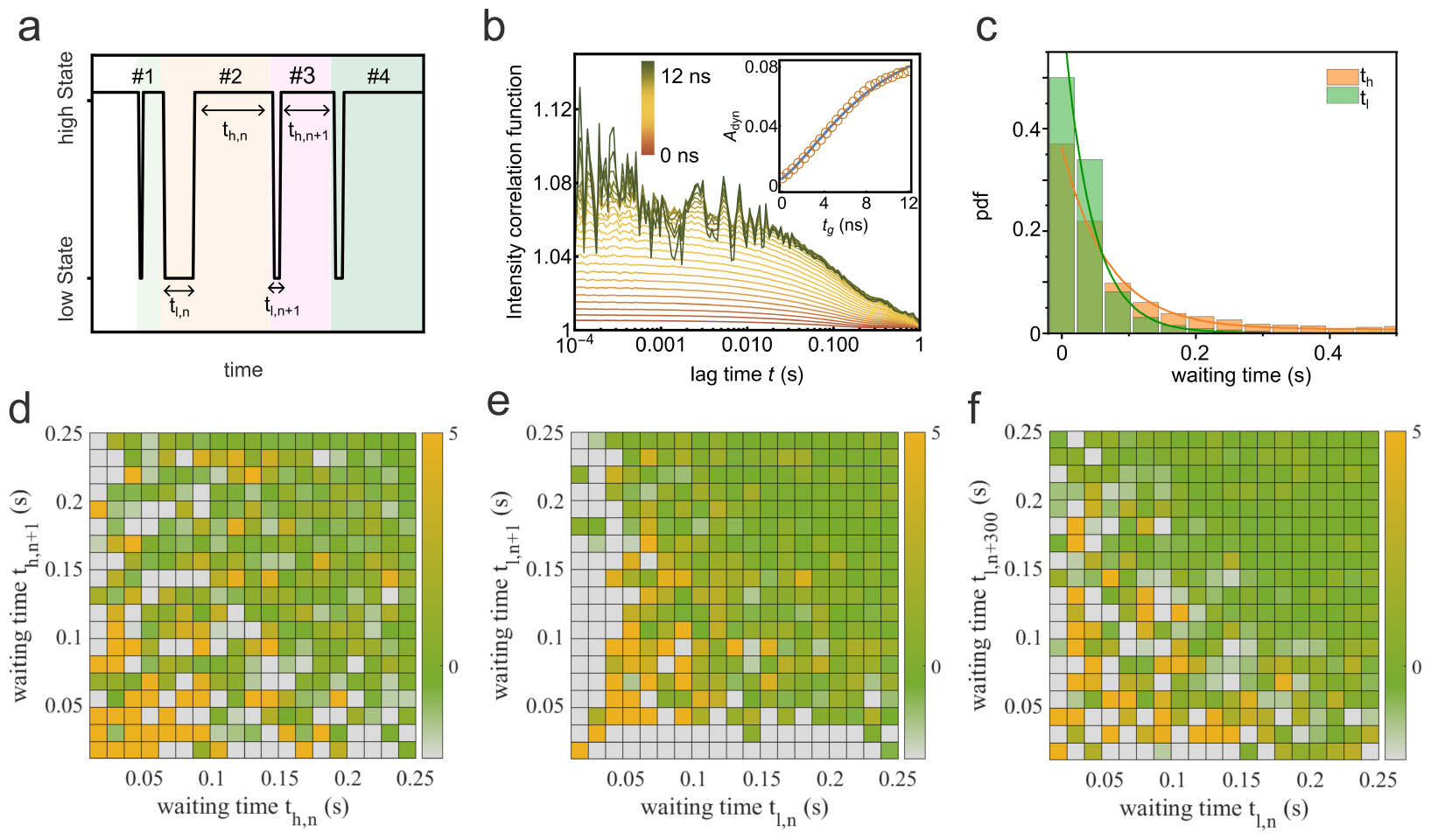
Waiting Time Correlation Analysis: (a) We define the waiting time for the low state as *t*_l_ and for the high state as t_h_ . These waiting times are calculated as the time intervals between two successive switching cycles. (b) The autocorrelation function is plotted as a function of the window width used for constructing the sg-FCS curves for the fluorescence intensity trace shown in Fig. 3a. The sg-FCS correlation functions are color-coded with a gradient from red to green, employing a 0.5 ns step size in increasing window width. These correlation functions were fitted with a mono-exponential decay function, and the resulting correlation amplitudes (A_dyn_ (t_g_)) are plotted in the panel inset. The data points are fitted with the model of eq. 6. (c) Possible distribution functions (pdf) of the waiting times t l and t h for more than 10,000 cycles. In (d) and (e), we present 2D difference histograms. These histograms illustrate the correlations between waiting times for consecutive cycles. Specifically, (d) shows the 2D difference histogram for t_h_, and (e) shows the same analysis for t_l_ . These plots reveal how waiting times correlate from one cycle to the next. In (f), we examine the 2D difference histogram of two waiting times for t_l_ but with a larger separation of 300 cycles lag. This analysis explores potential correlations over an extended period, providing insights into longer-term memory effects.

To dissect the formation dynamics of the E-cad X-dimer, we can individually describe each intensity time trace by the time spent in the high state (h) and low state (l) between consecutive height switching cycles, denoted as the waiting time for each state (t_i_, where i = h or l, as shown in Fig. 4a). These waiting times represent the duration required to transition to the next state. By averaging these waiting times across all intensity time traces, we obtain the mean waiting times for these two states: ⟨*t*_h_⟩ = 0.41 ± 1.29 s and ⟨*t*_*l*_⟩ = 0.046 ± 0.084 s. Here, ⟨· ⟩ denotes the averaging time, and errors are determined from the standard deviation of 13,160 switching cycles. The larger errors observed in waiting times might be attributed to the E-cads on each membrane not solely being monomers. X-dimers formed from the *cis*-dimers or *cis*-oligomers dramatically differ in stability. In addition, measurements using GUVs functionalized with a higher concentration of E-cad EC show a decrease in the average high state waiting time to 0.25 ± 0.56 s, while the low state waiting time does not vary significantly (0.051 ± 0.098 s, Supporting Information, Fig. S5). This suggests that the measured low state lifetime genuinely reflects the lifetime of the X-dimer.

Using these averaged waiting times, we can calculate the transition rate constants for switching between the low state and the high state: *k*_*l* → *h*_ = 1/ ⟨*t* _*l*_⟩ ≈ 22 s^− 1^ and from the high state to the low state: *k*_*l* → *h*_ = 1/ ⟨*t* _*h*_⟩ ≈ 2.4 *s*^− 1^ . This result s in an equilibrium constant (*K* = *k*_*h* → *l*_ / *k*_*l* → *h*_) of approximately0.11. Intriguingly, the rate constants for converting low state to high state is of the same order of magnitude as the rate constants for conformational switching between the S-dimer and X-dimer in solution, as measured by NMR relaxation dispersion spectroscopy (*k*_*X* → *S*_ = 86 s^− 1^) (13). This further suggests that the transitions between the high and low states in the fluorescence time trace primarily arise from conformational changes between dimers.

To provide further evidence of the dynamic changes between the high and low states, we employed a recently developed method called shrinking gate fluorescence correlation spectroscopy (sg-FCS) to determine the equilibrium constant (36). In sg-FCS, we generated a set of fluorescence intensity autocorrelation functions (iACFs) using distinct subsets of photons. This was achieved by adjusting the time window (on the nanoseconds time scale) following laser pulses, during which photons were considered for constructing the iACF. If dynamic changes are occurring, the amplitudes of the iACFs should incrementally increase with a longer time window.

As depicted in Fig. 4b, the iACFs for the membrane with E-cad clearly exhibit an increase in amplitudes as the time window is shrunk, with a step size of 0.5 ns. By fitting the curve amplitudes against window width, we determined the equilibrium constant to be approximately 0.1. Interestingly, this equilibrium constant obtained using sg-FCS closely matches the K value calculated from the waiting times. This agreement strengthens the conclusion that the dynamic transitions between high and low states primarily result from conformational changes between dimers.

The distribution of individual waiting times follows an exponential decay function for both *t*_*h*_ and *t*_*l*_, both displaying broad distributions (Fig. 4c). Further, a typical trajectory in Fig. S8 illustrates the stochastic appearance of waiting times for each state. This raises the question of whether these fluctuations in waiting times for each state exhibit temporal correlations. In other words, can the appearance of two consecutive waiting times be statistically correlated with temporal changes in the structures of the dimers?

To investigate these potential correlations, we evaluated the 2D difference histogram, defined as *δ*(*t*_*i, n*_, *t* _*i, n* + 1_) = *p*(*t*_*i, n*_, *t* _*i, n* + 1_) − *p*(*t*_*i, n*_) × *p*(*t*_*i, n* + 1_), for the waiting times of each state from two adjacent cycles (see Supporting Information). This analysis involves calculating the joint probability for two waiting times, denoted as *p*(*t*_*i, n*_, *t* _*i, n* + *1*_), and comparing it to the probability of no correlation between these two times, which is represented by the product of the individual waiting time probabilities, *p*(*t*_*i, n*_) × *p*(*t*_*i, n* + 1_). This statistical t est has previously been employed ins the analysis of dynamic disorder in enzymatic catalysis (37–39).

In this context, if there is no dynamic disorder, the joint probability *p*(*t*_*i, n*_, *t* _*i, n* + 1_) and the product of individualprobabilities *p*(*t*_*i, n*_) × *p*(*t*_*i, n* + 1_) would be the same (see Supporting Information). If they are different, it indicates the presence of a memory effect induced by conformational changes, where the system ‘remembers’ its conformation for an extended period.

We conducted a thorough analysis of waiting times from all analyzed trajectories (N = 25, each spanning a 600 s experiment). Notably, we observed distinct waiting time distributions for the high and low states.

As depicted in Fig. 4c and Fig. 4d, the waiting times for the high state did not exhibit significant time-correlated features. In contrast, for the low state, we identified a weak and broad diagonal feature in the waiting time distributions. This diagonal feature suggests that longer waiting times are less likely to be followed by shorter ones and vice versa. This intriguing pattern implies a form of temporal memory within the system.

Remarkably, this correlation in waiting times for the low state persisted even when considering cycle lags of 20, corresponding to a time range on the order of seconds (see Supporting Information, Fig. S10). Such memory effects have been previously observed in enzymatic catalysis and attributed to conformational fluctuations in individual enzymes (37–39).

However, in our experimental context, the low states result from the formation of numerous X-dimers, with E-cad molecular densities estimated to be approximately 100-700 molecules per square micrometer within each membrane (24). Consequently, the observed correlation or memory effect in the low state is less likely to stem from the conformational dynamics of individual X-dimers. Instead, it may arise from inter-dimer communication among the E-cad dimers. Specifically, the presence of X-dimeric states can influence the structure of nearby S-dimers by altering the membrane-membrane distance, potentially affecting the stability and persistence of X-dimers formed from these S-dimers.

## Discussion

We employed MIET spectroscopy to probe the intricate membrane-membrane adhesion process facilitated by Ecadherins (E-cads) on each membrane. Our investigations centered on a biomimetic system designed to emulate E-cad-mediated cell adhesion. Specifically, we explored the adhesion events of the EC domains of E-cads, which were positioned between supported lipid bilayers (SLBs) and giant unilamellar vesicles (GUVs).

Consistent with earlier reports highlighting the all-or-nothing nature of E-cad junction formation, our observations revealed two markedly distinct adhesion conformations.

In the first scenario, we observed a conventional E-cad-mediated adhesion process, resulting in the formation of a *cis*-cluster between the two membranes. Notably, we observed variations in membrane height during this adhesion process. Leveraging MIET, we precisely determined the height values at different stages of this process. These height values, coupled with crystal structure parameters for E-cad and E-cad junctions, led us to conclude that E-cad experiences a transition from an initial dimeric state with a larger EC5-EC5 distance to reach a final, stable *cis*-clustered state with a smaller distance.

In contrast, the second scenario involved the observation of only the dimeric state without further clustering. Intriguingly, we detected continuous ‘dip’ signals in the fluorescence intensity trace. Through analysis of the EC5-EC5 distance derived from these ‘dip’ signals, our data strongly support the hypothesis of an X-dimer-dependent mechanism for wild-type E-cad mediated membrane adhesion, with the ‘dips’ indicative of X-dimer formation. Furthermore, we uncovered the kinetics of X-dimer formation and dissociation, revealing that the dissociation rate of X-dimer is tenfold faster than the formation rate. The X-dimer exhibited a brief lifetime of approximately 50 ms, underscoring its role as a short-lived intermediate state. Our MIET -based analysis extended to quantifying membrane fluctuation amplitudes. We discovered that the conformational dynamics of dimers lead to more pronounced membrane displacements. Interestingly, these membrane displacements triggered a ‘memory effect’ among X-dimer formations, hinting at mutual cooperativity between membrane displacement and trans-dimers. While prior simulations and experiments have underscored mutual *cis*/ *trans* cooperativity in E-cad, stemming from apparent allosteric enhancement, our results suggest a novel mechanism whereby membrane displacement also contributes to this cooperativity.

In summary, our utilization of MIET spectroscopy and imaging enabled us to monitor the height variations in a model E-cad-modified membrane system. We delved into the kinetic and dynamic processes underlying E-cad-mediated membrane adhesion, pinpointing various E-cad states based on their EC5-EC5 distances and studying their evolution during adhesion. Notably, we unraveled the existence of an X-dimeric intermediate state characterized by its transient nature and its significant impact on membrane displacement, which, in turn, influences the formation and stability of subsequent X-dimers. The ability of X-dimers to communicate with one another through membrane displacement further underscores the intricate mechanisms underlying junction dynamics. These insights are crucial for understanding the role of E-cad junctions in maintaining cell-cell adhesion, tissue organization, and physiological processes. Further, our findings underscore the potential of MIET spectroscopy and imaging as a potent technique for elucidating subtle structural changes in membrane proteins.

## Materials and Methods

Details of the materials (lipids and proteins) are available in SI Materials and Methods.

### Substrate preparation

The gold-coated coverslips were prepared with the atom vapor deposition method. Briefly, commercial coverslips (d = 150 μm) were cleaned by the following detergent treatment : (1) soaking in the piranha lotion for 30 min, (2) ultrasonication in 5% KOH for 15 min, (3) ultrasonication in ethanol for 15 min, and (4) ultrasonication in DI water for 15 min. Steps 2 to 4 were repeated twice. The cleaned coverslips were further used as substrates for vapor deposition of gold and SiO_2_ spacer. Under high-vacuum conditions, 2 nm titanium, 10 nm gold, 1 nm titanium, and 10 nm SiO_2_ were evaporated on the surface of the coverslip layer by layer by using an electron beam source. The lowest deposition rate was maintained at 1 Å s^− 1^ to ensure maximum smoothness on the surface.

### Supported lipid bilayer (SLB)

SLBs were prepared with the Langmuir-Blodgett Langmuir-Schäfer (LB-LS) techniques with a Langmuir-Blodgett trough (Nima, Coventry, UK). The subphase was ultrapure water. The proximal monolayer consisted of pure SOPC or DPPC and the distal layer was formed by SOPC or DPPC with 2 mol% PEG2000-DOPE, and 5 mol% DOGS-NTA . For DPPC SLB, 1% NBD-PC was added to the distal layer. The transfer pressure was maintained at 28 mM / m for SOPC and 30 mM / m for DPPC. The SLBs were kept wet all the time and used directly after preparation.

### Giant unilamellar vesicles (GUV)

GUV was prepared by the electroswelling method. Lipids chloroform mixture consisting of SOPC with 2 mol% PEG2000-DOPE, 1 mol% Atto655-DPPE, and 5 mol% DOGS-NTA was deposited on a home-build electrode and evaporated for 3 h under vacuum at 30°C. The electrode was assembled into a chamber and filled with 500 μL of 230 mM sucrose solution. Then an alternation current at 15 Hz for 3 h and a peak-to-peak voltage of 1.6 followed by 8 Hz for 30 min were applied to the chamber. The GUVs suspension was st ored at 4°C and used within 3 days.

### Measurement sample preparation

For preparing the measurement sample, SLB was first incubated with 2 μM NiSO_4_ PBS solution for 15 min and then washed with PBS 5 times. Afterward, This SLB was further exposed to 10 μg/ mL E-cad PBS solution for 3 h. Meanwhile, the stock GUVs solution was also incubated with the same concentrations of NiSO_4_ and E-cad for 3 h. Then the E-cad-functionalized GUVs were diluted 100 times with PBS-CaCl_2_ buffer (140 mM NaCl, 3 mM KCl, 10 mM Na_2_HPO_4_, 2 mM K H_2_PO_4_,750 μM CaCl_2_, pH 7.4). Finally, the diluted GUVs solution was added into the chamber with SLBs, and measurements were started immediately. To avoid solution evaporation, a glass slide was used to cover the chamber during the measurement .

### MIET measurement

All measurements were carried out with homebuilt confocal microscopy equipped with a time-correlated single photon counting (TCSPC) device. A pulsed diode laser (λ_ex c_= 640nm, LDH-D-C 640, PicoQuant), which has a pulse width of 50 ps FWHM and repetition rate of 40 MHz, and a clean-up filter (LD01-640/ 8 Semrock) worked as the excitation source. A beam of 12 mm diameter, which was collimated from an infinity-corrected 4x objective (UPISApo 4X, Olympus), was reflected by a dichroic mirror (Di01-R405/ 488/ 561/ 635, Semrock) t owards a high numericalaperture objective (UApoN 100X, oil, 1.49 N.A ., Olympus). The emission light was focused into a pinhole of 100 μm diameter and refocused onto two avalanche photodiodes (τ - SPA D, PicoQuant). Additionally, a long-pass filter (BLP01-647R-25, Semrock) and two band-pass filters (Brightline HC692/ 40, Semrock) were used before the pinhole and the detectors, separately. Photo signals acquired from the detector were processed by a multi-channel picosecond event timer (Hydraharp 400, PicoQuant) with 16 ps time resolution. In a typical measurement, we scanned the sample at the coverslip surface to determine the contour of the GUV and choose the appropriate point position for data acquisition. The power of the laser is adjusted until a maximum count rate of 100-500 kcps is reached.

### Principle of MIET

The theory for converting the lifetime values to the substrate-fluorophore distances with MIET has been elaborated in detail in several previous publications(31–33). Briefly, the electromagnetic field of a fluorescing molecule (electric dipole emitter) is represented by a superposition of plane electromagnetic waves. When interacting with the MIET substrate, each plane wave is reflected and transmitted based on Fresnel’s reflection and transmission laws. The full field is Thus represented by a superposition of the incident and reflected (molecule’s half space) or transmitted plane waves (objective’s half space). The emission rat e of the fluorescent molecule is then calculated by integrating the Poynting vector of the full field over a closed surface enclosing the emitter. Thus, we get an expression for the radiative emission power (S(θ, z_0_)) which can be decomposed as

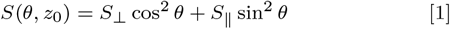

where z_0_ is the distance of the emitter from the surface, θ is the orientation angle of This emitter (angle between emission dipole axis and vertical axis), and *S*_⊥_ (or *S*_•_) are the radiative emission rates of a dipole emitter orientated perpendicular or parallel to the subst rat e, respectively. Taking the molecules’ nonradiative transition into account, the expression for the fluorescence lifetime (*τ*_f_ (*θ, z*_0_)) of a molecule is given by

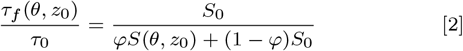

where *τ*_0_ is the lifetime of the molecule in free space without any MIET substrate, *ϕ* is the quantum yield of the molecule, and S_0_ is the emission power the emitter in free space. Furthermore, the relative brightness *b*(*θ, z*_0_) of an emitter near the metal surface is proportional to the product of the position- and orient ation-dependent collection efficiency *η*(*θ, z*_0_) of fluorescence det ection and the local quantum yield *ϕ* _l ocal_ (*θ, z*_0_). The former is found by integrating the position- and orientation-dependent angular distribution of emission of the emitter over the cone of light collection of the microscope’s objective. The local quantum yield *ϕ*_*local*_ (*θ, z*_0_) is defined by

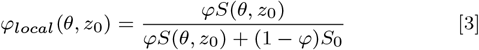

Thus, we find

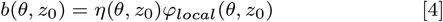

Fig. 2c shows the calculated lifetime and relative brightness as a function of distance z for the case of an emitter oriented randomly to the substrate. For the DPPE-Atto655 labeled membrane, the free space lifetime was measured on a GUV sample on a glass surface without gold film, and the quantum yield (*ϕ* = 0.36) was measured using a nanocavity method (40). Note that when the membrane forms the final *cis*-cluster state, the membrane does almost not fluctuate. In this case, we assumed a dye orientation parallel to the substrate surface when calculating the calibration curve (32).

A custom-written MATLAB-based software package for the calculation of MIET lifetime-versus-distance curves as well as the conversion of lifetime to distance, equipped with a graphical user interface, has been published (32) and is available free of charge at https://projects.gwdg.de/projects/miet.

### Conversion of intensity to height correlation curves

In general, the autocorrelation function for conventional FCS is defined as while the height correlation function (g_h_) for the membrane fluct u-ations is defined as

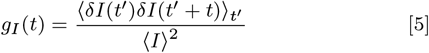

while the height correlation function (*g*_*h*_) for the membrane fluctuations is defined as (*g*_*h*_ (*t*) = .*rδh*(*t* ^*M*^)*δh*(*t* ^*M*^ + *t*)._*t M*_, where *δh*(*t*) is the membrane displacement from its average position.

The relationship between fluorescence intensity (*I*) and membrane height (*h*) can be determined from eq. 4. However, to simplify the conversion, the *I* − *h* relationship can be estimated approximately from the experiment . Fig. S6 shows a two-dimensional histogram of the fluorescence intensity and fitted lifetimes measured from the sample without E-cad between GUV and SLB, it can be found that the dependence of intensity on height is almost linear. Thus we take a linear fitting on the counts versus height s dat a, which gives the slope (*m* = *δI* / *δh*). Finally, we obtained the *g*_*h*_ (*t*) from the conversion of *g*_*I*_ (*t*):

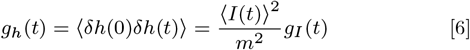

### sg-FCS analysis

The theory for analyzing the sg-FCS has been thoroughly elaborated by Schröder et al. (36). In brief, each photon is recorded with two-time tags, namely the macro-time and microtime, which respectively refer to the photon’s arrival time from the beginning of the experiment and the delay time between the laser pulse and the photon detection. We generated a series of sg-FCS curves (iACFs) using different subsets of photons depending on their micro-time. Then, the bunching amplitude (*A*_*dyn*_) for each iACF was obtained by fitting a mono-exponential model. At last, we determined the equilibrium constant (*K*) by fitting the correlation amplitude versus sg-FCS time window width using the following model equation

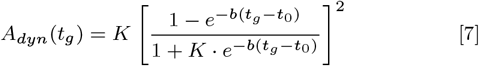

where *t*_0_ is the temporal position of the laser excitation pulsepeak, and *b* denot es an energy transfer rate constant . It ‘s important to note that all iACFs were background-corrected by measuring the background signal on an empty gold-coated coverslip.

### Data and program availability

All data of the figures in This manuscript together with Matlab and Mathematica programs that were used to generate these figures can be accessed here: https://gitlab.gwdg.de/igregor/e-cadherin.git.

## ACKNOWLEDGMENTS

J.E. acknowledges financial support by the DFG through Germany’s Excellence Strategy EXC 2067/ 1-390729940. T .C. and J.E. thank the European Research Council (ERC) for financial support via project “smM IET” (grant agreement no. 884488) under the European Union’s Horizon 2020 research and innovation program.

