## Supporting Information for "Observation of E-cadherin Adherens Junction Dynamics with Metal-Induced Energy Transfer Imaging and Spectroscopy"

#### This PDF file includes:

Supporting text
Figs. S1 to S10
Legends for Movies S1 to S2
SI References

#### Other supporting materials for this manuscript include the following:

Movies S1 to S2

### Supporting Information Text

**Materials.** Lipids, 1-stearoyl-2-oleoyl-sn-glycero-3-phosphocholine (SOPC), 1,2-dipalmitoyl-sn-glycero-3-phosphocholine (DPPC), 1,2-dioleoyl-sn-glycero-3-[(N-(5-amino-1-carboxypentyl)iminodiacetic acid)succinyl] (nickel salt) (DOGS), 1,2-distearoyl-sn-glycero-3-phosphoethanolamine-N-[methoxy(polyethylene glycol)-2000] (ammonium salt) (PEG2000-DOPE), 1,2-dioleoyl-sn-glycero-3-phosphoethanolamine-N-(cap biotinyl) (sodium salt) (DOPE-cap-Biotin), and 1-palmitoyl-2-6-[(7-nitro-2-1,3-benzoxadiazol-4-yl)amino]hexanoyl-sn-glycero-3-phosphocholine (NBD-PC) were purchased from Avanti Polar Lipids and diluted at 10 mg/mL with chloroform for stock solution. Atto655 labeled 1,2-Dipalmitoyl-sn-glycero-3-phosphoethanolamine (DPPE) (Atto655-DPPE) was purchased from Atto-TEC and dissolved in chloroform at 1 mg/mL for stock solution.

Proteins, neutravidin, and E-Cadherin-Fc chimera were used following the manufacturer's instructions. The neutravidin protein (Thermo ScientificTM) was reconstituted with ultrapure water and then diluted to 1 mg/mL with PBS (140 mM NaCl, 3 mM KCl, 10 mM $Na_2HPO_4$, 2mM $KH_2PO_4$, pH 7.4). The recombinant human E-cadherin-Fc chimera (referred to as E-cad, Biolegend) was reconstituted with PBS at 100 $\mu$g/mL and stored at -80°C.

**Suppression of diffusion signal in FCS by using high dye concentration.** To mitigate the influence of lipid diffusion on our fluorescence intensity-dependent measurements, we employed a high dye concentration (1 mol%) of DPPE-atto655 in the lipid bilayer. This strategy, recommended by Monzel et al. (1), effectively minimizes diffusion-related signals. To confirm that this dye concentration adequately eliminates fluctuations induced by diffusion, we conducted fluorescence correlation spectroscopy (FCS) on supported lipid bilayers (SLBs) with varying dye concentrations.

In Figure S1a, the autocorrelation function (ACF) obtained from an SLB with a dye concentration of 0.001 mol% reveals modulation in the correlation curve due to the diffusion of a limited number of fluorophores in the confocal volume. In contrast, when the dye concentration in the SLB is increased to 1 mol% (Figure S1b, black curve), the FCS signal diminishes, and the ACF flattens, indicating effective suppression of diffusion-related noise.

Furthermore, we performed FCS on a fluctuating membrane by immersing a highly labeled GUV (1 mol%) in a hypertonic solution (300 mOsm $l^{-1}$, PBS). In this scenario, the GUV becomes floppy, and its membrane undergoes significant bending fluctuations. As anticipated, we observed the ACF for such a deflated GUV (Fig. S1b red curve). Due to the high dye labeling, this signal is devoid of any diffusion contribution, and the amplitude solely results from physical displacements of the membrane.

**Fluorescence Recovery after Photobleaching (FRAP) measurements.** To investigate the diffusion dynamics of NBD-PC-labeled lipids in both solid-supported phospholipid bilayers (SLBs) of 1-palmitoyl-2-oleoyl-sn-glycero-3-phosphocholine (SOPC) and 1,2-dioleoyl-sn-glycero-3-phosphocholine (DPPC), fluorescence recovery after photobleaching (FRAP) measurements were employed. Both SOPC and DPPC SLBs were prepared using the Langmuir-Blodgett (LB)-Langmuir-Schaefer (LS) method, with 1 mol% NBD-PC, 2 mol% polyethylene glycol (PEG) 2000-doped DOPE, and 5 mol% DOGS-NTA incorporated into the distal layer of the bilayer. A 488 nm laser beam was used to photobleach the NBD-PC in the focus area using the highest laser power for 1 second, achieving an average bleach area radius of 3.75 micrometers. Fluorescence images were captured immediately after photobleaching.

In SOPC measurements, owing to the swift diffusion of NBD-PC, continuous fluorescence imaging was executed without interruption at a rate of 250 nanometers per 10 microseconds (pixel size/dwell time) for an area of 50 by 50 micrometers squared. For DPPC measurements, fluorescence images were captured every 2 minutes over a span of at least 40 minutes. The fluorescence intensity of the bleached spot was normalized to the fluorescence intensity of an unbleached reference spot, and the relative fluorescence intensity was plotted over time.

Figure S3 illustrates a representative FRAP recovery profile of fluorescence-labeled SOPC SLB. The red curve depicts a single exponential fit to the data using the following equation:

$$y = A(1 - e^{-bx}) \tag{1}$$

where $A$ corresponds to the mobile fraction and $b$ is related to the diffusion time via

$$t_{1/2} = \frac{\ln 2}{b} \tag{2}$$

which is related to the diffusion constant by

$$D = \frac{w^2}{4t_{1/2}} \tag{3}$$

where $w$ is the radius of the bleaching area. Thus, we obtained a diffusion constant 0.5 $\mu m^2/s$ for SOPC with NBD-PC SLB and of 0.004 $\mu m^2/s$ for DPPC with NBD-PC. Note that the value of $D$ for SOPC is much smaller than the previously reported value for a fluid SOPC bilayer which shows a value of $D$ larger than 1 $\mu m^2/s$ (2). This is due to the fact that in our experiments, we did not take into account the overall photobleaching arising from the continuous scanning during the SOPC measurement.

**Membrane mobility and E-cad mediated intermembrane adhesion.** Fig. S4a and Fig. S4b show two typical time traces measured from E-cad decorated GUVs on E-cad decorated fluid SOPC SLBs. The clustering timescales (blue regions) for the fluid SOPC SLBs are in the range of 1.2-1.4 s, While for the measurement of partially fluid SLBs made of DPPC and NBD-PC (Fig. S4c), the clustering timescales range from 15 s to 30 s. This observation indicates that the *cis*-clustering of E-cad is strongly dependent on membrane mobility.

**Confirmation of the origin of consecutive 'dip signals' .** Having recognized that the dips in intensity result from the appearance of the X-dimer within the confocal volume, a question arises regarding whether the consecutive 'dip signals' originate from the formation of X-dimer within the confocal volume or the diffusion of X-dimer. This diffusion could involve the X-dimer forming outside the confocal volume and subsequently diffusing in and out of it. To verify this, we measured the diffusion coefficient of the SLB prepared using DPPC and NBD-PC.

Previous reports indicate that these partially fluid SLB membranes, not in a full gel phase at room temperature, are not expected to be phase-separated due to the low density of Ni-NTA-DOGS used (2, 3). The diffusion coefficient of NBD-PC lipids in this SLB is estimated to be 0.004 $\mu$m$^2$/s from fluorescence recovery after photobleaching (FRAP) experiments (Fig. S3), corresponding to a diffusion time of approximately 5 s. This implies that the 'dips' are definitely not caused by diffusion, as the average waiting time of 'dip signals' is approximately 0.05 s, much shorter than the diffusion time.

**Determination of the high and low states.** Due to thermal fluctuations in the membrane, the high state in the time trace exhibits significant noise, leading to a low signal-to-noise (SN) level. In Fig. S7b, a distribution histogram of count rates from a typical time trace reveals the challenge of completely separating the two states using a single dividing line, given the overlap between them.

To identify the high and low states, we employed a two-threshold strategy for isolating the low state from the noisy high states. In this approach, a high-threshold (proximate to the high state, Fig. S7a red line) was established to select all bursts with intensities surpassing the threshold. Subsequently, for each selected low state, the total fluorescence counts were integrated, and a low threshold was set to filter out bursts with intensities exceeding the high-threshold (Fig. S7a blue line). This two-threshold strategy ensures the inclusion of some low states with high fluorescence intensity that persist for an extended duration in the time trace.

Firstly, part of the histogram of the intensity that definitely has no low states was fitted with a Gaussian distribution:

$$I = A \exp\left[\frac{-(I - I_0)^2}{2\sigma_1^2}\right] \quad [4]$$

where $I$ is the intensity in counts per time bin, $I_0$ is the center of the Gaussian distribution, and $\sigma_1$ is the standard deviation of the photon count distribution.

Secondly, the time traces $I(t)$ were shifted to zero mean value by $I(t) - I_0$. Then, the variance $\sigma_1$ was used to set the thresholds for determining the low states and high states.

We chose the low states from the control experiment (GUV without E-cad) as $N_{noise}$. The number of low states extracted from the intensity trace of E-cad binding was designated as $N_{signal}$. We then calculated $N_{noise}$ and $N_{signal}$ based on the two-threshold method by setting different threshold values in units of $\sigma_1(i \times \sigma_1)$ for these two thresholds.

Next, we calculated the ratio $N_{noise}/N_{signal}$ as function of the values of the low and high thresholds. Fig. S7c shows the dependence of $N_{noise}/N_{signal}$ on the combination of the two thresholds. The red and yellow lines indicate the ratio of 10% and 5%, respectively. We found that when setting the high state at $2.5\sigma_1$ and the low state at $3\sigma_1$, the $N_{noise}/N_{signal}$ is smaller than 2%, which means only 2% of the signal from the high state is counted as coming from the low states. We thus chose $2.5\sigma_1$ and $3\sigma_1$ as the high and low thresholds for separating the low states and high states in the fluorescence intensity trace.

**Construction of the 2D difference histograms of waiting times.** A 2D conditional histogram of waiting times for the high or low state describes the joint probability of finding pairs of waiting times ($t_{i,n}$ and $t_{i,n+1}$, where i = h or l). We also generated a correlation plot between the waiting time $t_{i,n}$ and $t_{i,n+300}$ (Fig. S9). At this larger separation, the correlation has diminished.

For the waiting time from the low state ($t_l$), the 2D joint histograms between the lag of 1 'low state' time and the lag of 300 'low state' times exhibit substantial changes: there is a notable excess of points distributed along the axes at larger lags ($t_{l,n}$ vs. $t_{l,n+300}$), while the scatter plots follow the inverse diagonals for two adjacent waiting times ($t_{l,n}$ vs. $t_{l,n+1}$). Conversely, for the waiting time from the high state ($t_h$), almost no difference was observed. The two histograms between the low states with different lags indicate a significant difference, suggesting the presence of a memory effect.

To accentuate the memory effect, 2-D difference histograms of waiting times were constructed. The 2-D difference histograms are defined as $\sigma(t_{i,n}, t_{i,n+j}) = p(t_{i,n}, t_{i,n+j}) - p(t_{i,n}) \times p(t_{i,n+j})$ for correlation analysis of one state (high state or low state), where j represents the number of cycles that separate waiting time $t_n$ and $t_{n+j}$.

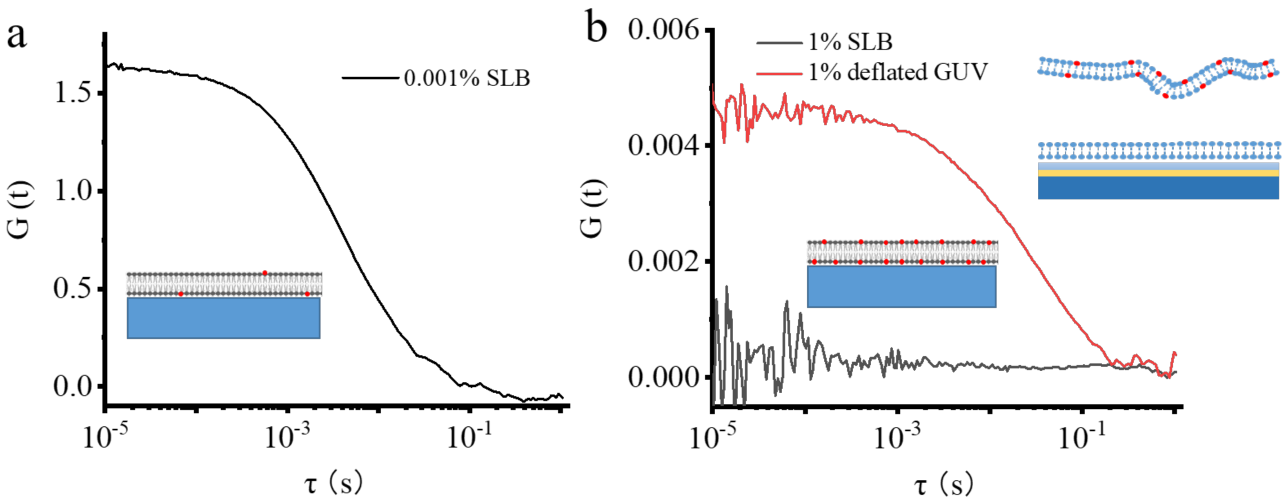


**Fig. S1.** ACFs measured on (a) lowly labeled SLB, (b) highly labeled SLB (black curve), and highly labeled deflated GUV (red curve).

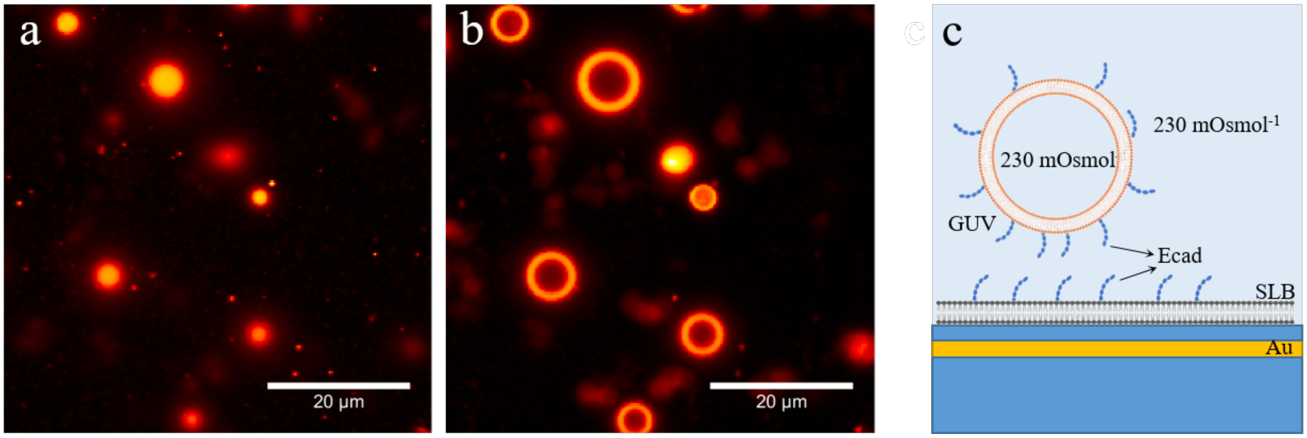


**Fig. S2.** Fluorescence images of E-cad-modified GUV on E-cad-modified SLB recorded (a) directly on the surface and (b) at a height of 1 $\mu$m above the surface. (c) Schematic representation of the E-cad-modified tense GUV on E-cad-modified SLB, above a gold surface with 10 nm silica spacer. Tense GUVs were prepared by immersing them into a PBS solution (230 mOsmol$^{-1}$), which matches the osmolality of the sucrose solution (230 mOsmol$^{-1}$) inside the GUV.

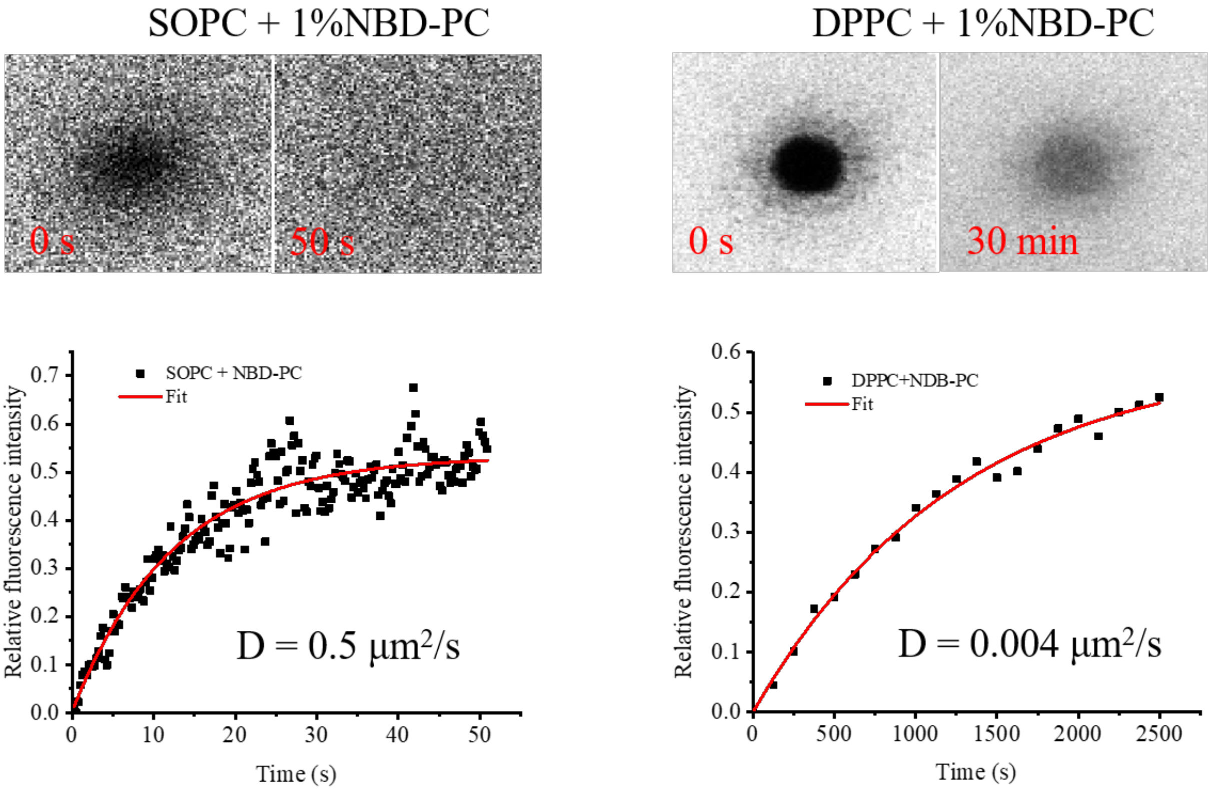


**Fig. S3.** FRAP analysis of SOPC (a) and DPPC (b) SLBs. Top: photobleached spot that recovers after 50 s (a) and 30 min (b). Bottom: fluorescence recovery profile after photobleaching.

 

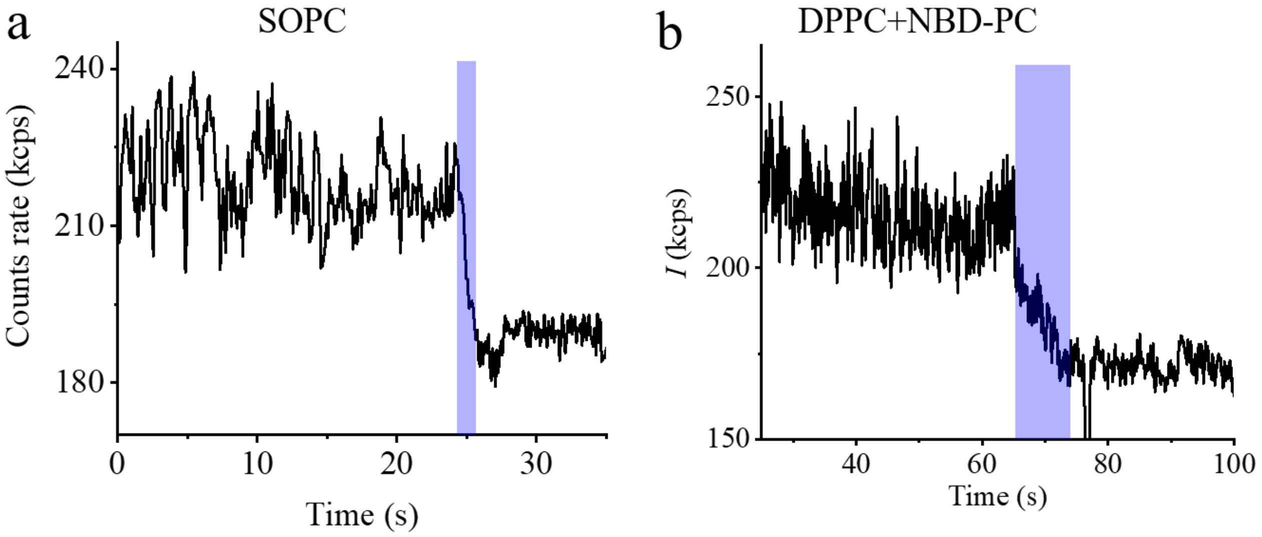


**Fig. S4.** Two typical fluorescence time traces of E-cad-mediated membrane adhesion with the formation of the final cic-cluster state. The trace was measured from the biomimetic system on (a) fluid SOPC SLB and (b) partially fluid SLB made of DPPC and NBD-PC.

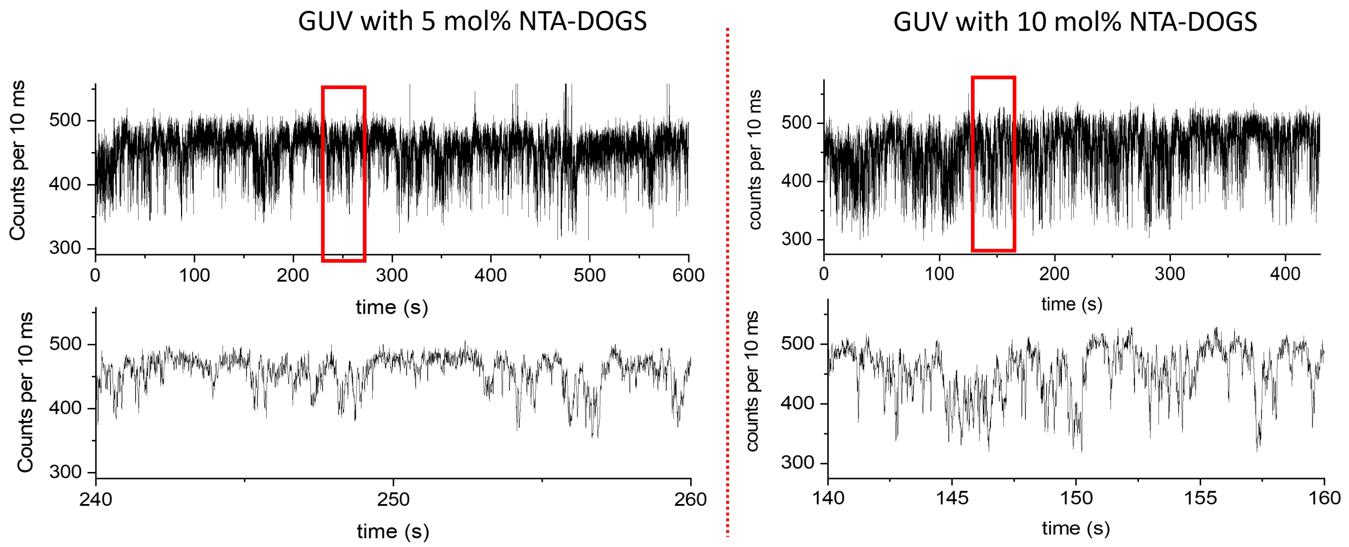


**Fig. S5.** Fluorescence intensity time traces obtained from GUVs with two different concentrations of E-cadherin (Ecad) on partially fluid SLBs. Enlarged plots are shown at the bottom.

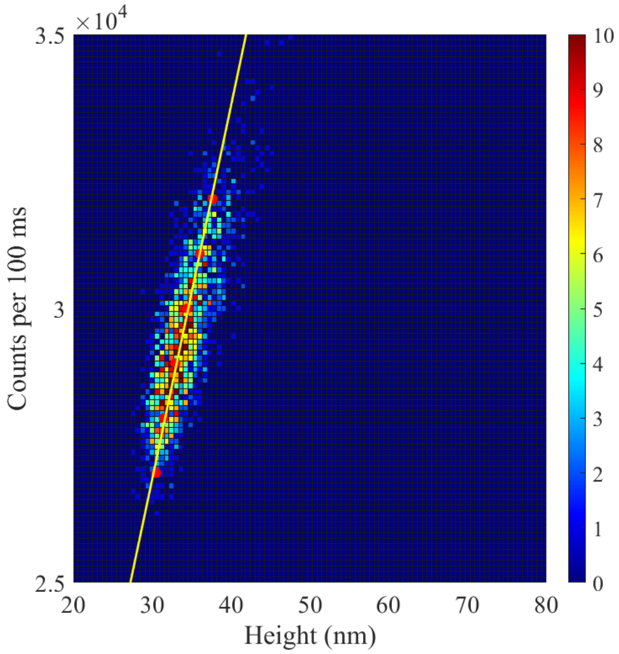


**Fig. S6.** A two-dimensional histogram of the fluorescence intensity and fitted lifetimes, data was measured on a sample without E-cad between GUV and SLB. The yellow line is a linear fit.

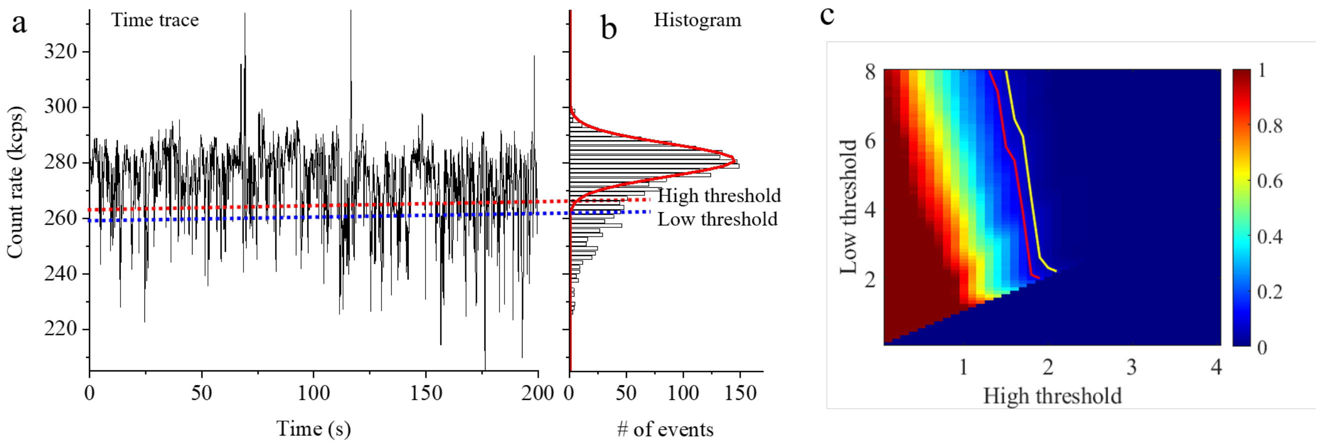


**Fig. S7.** (a) Fluorescence intensity trace and (b) corresponding intensity histogram. (c, d) Dependence of Nnoise/Nsignal on the combination of high and low threshold.

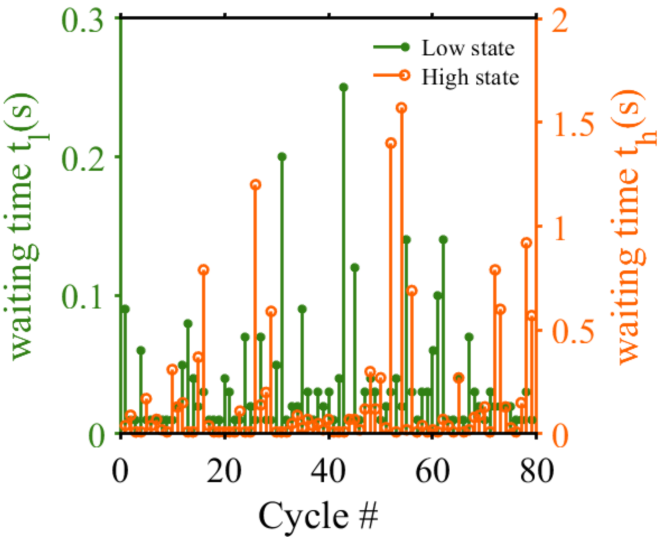


**Fig. S8.** Waiting time trajectories for $t_l$ and $t_h$ reconstructed from fluorescence time trace.

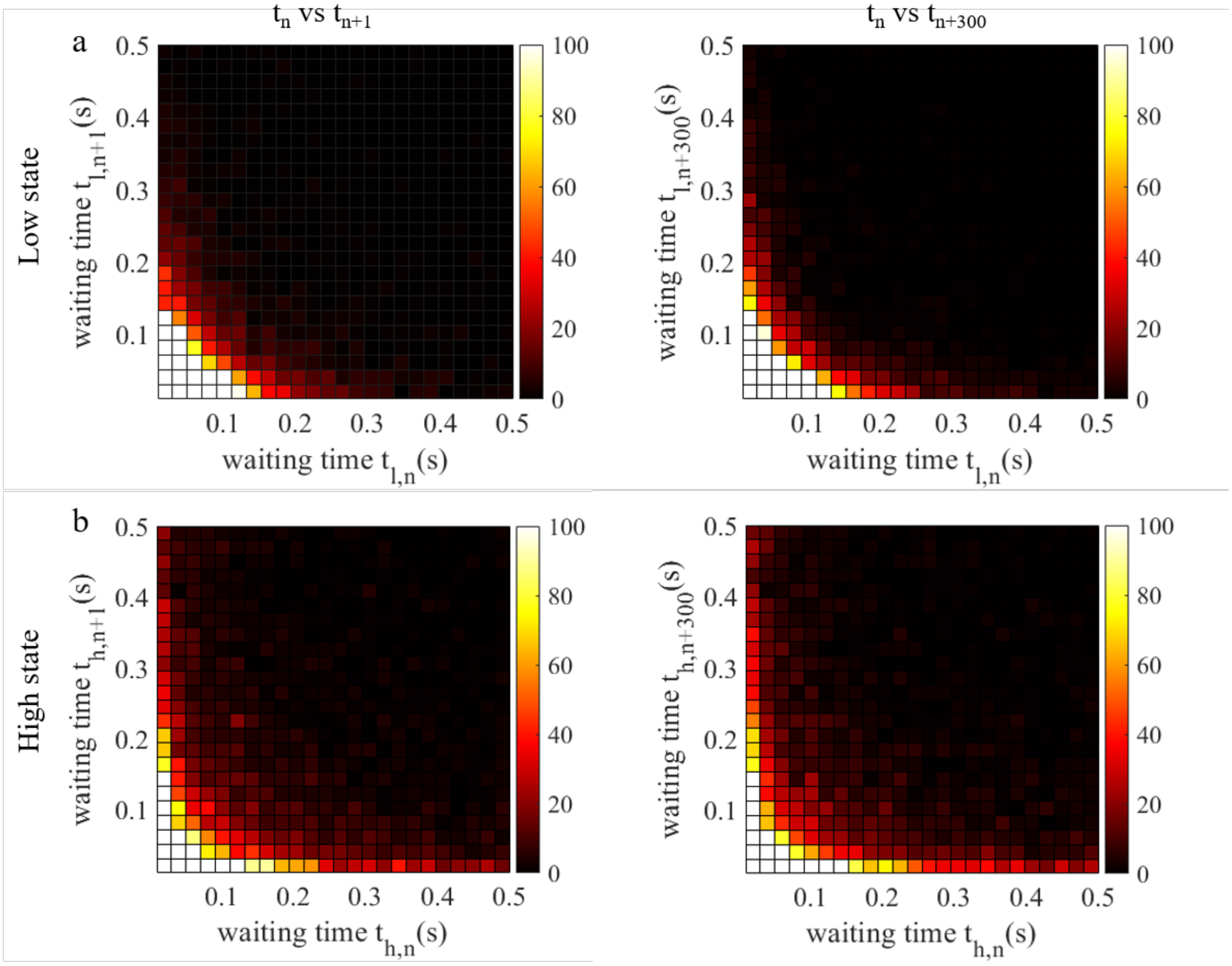


**Fig. S9.** 2D conditional histograms of waiting times for low state (a) and high state (b) recorded at $t_n$ and $t_{n+1}$ (left panel) as well as $t_n$ and $t_{n+300}$ (right panel).

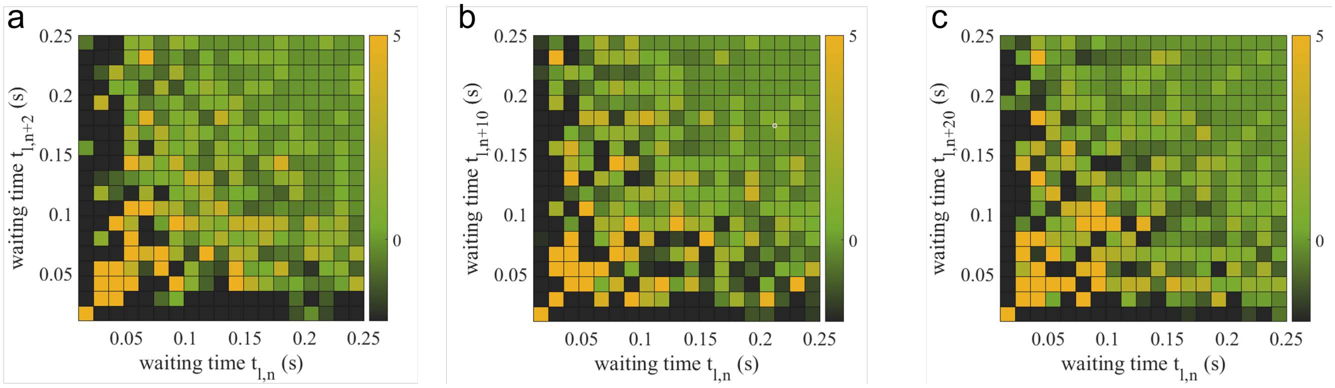


**Fig. S10.** 2D difference histograms of waiting times for low state, calculated for (a) $t_n$ and $t_{n+2}$, (b) $t_n$ and $t_{n+10}$, and (c) $t_n$ and $t_{n+20}$.

**Movie S1. This movie shows a process of E-cad-modified GUV binds to an E-cad-modified SLB and finally adheres to the SLB due to the formation of the *cis*-clustered state of the E-cads.**

**Movie S2. The movie shows the fluctuation of an E-cad-modified GUV above an E-cad-modified SLB. Notably, adhesion does not occur due to the absence of the *cis*-clustered state formation.**
